# Enhancing fMRI decoded neurofeedback with co-adaptive training: simulation and proof-of-principle evidence

**DOI:** 10.1101/2025.02.21.639408

**Authors:** Najemeddine Abdennour, Pedro Margolles, David Soto

**Affiliations:** Basque Center on Cognition, Brain and Language, Paseo Mikeletegi 69, 2nd Floor 20009 San Sebastian

**Keywords:** real-time functional magnetic resonance imaging, co-adaptation, decoded neurofeedback, machine learning

## Abstract

A significant challenge for neurofeedback training research and related clinical applications, is participants’ difficulty in learning to induce specific brain patterns during training. Here, we address this issue in the context of fMRI-based decoded neurofeedback (DecNef). Arguably, discrepancies between the data used to construct the decoder and the data used for neurofeedback training, such as differences in data distributions and experimental contexts, neural and non-neural noise, are likely the cause of the difficulties of the aforementioned participants. Here, we developed a co-adaptation procedure using standard machine learning algorithms. The procedure involves an adaptive decoder algorithm that is updated in real time based on its predictions across neurofeedback trials. First, we tested the procedure via simulations using a previous DecNef dataset and showed that decoder co-adaptation can improve performance during neurofeedback training. Importantly, a drift analysis demonstrated the stability of the co-adapted decoder throughout the neurofeedback training sessions. We then collected real time fMRI data in a DecNef training procedure to provide proof of concept evidence that co-adaptation enhances participant’s ability to induce the target state during training. Thus, personalized decoders through co-adaptation can improve the precision and reliability of DecNef training protocols to target specific brain representations, with ramifications in translational research. The tools are made openly available to the scientific community.

## I. Introduction

Decoded neurofeedback (DecNef) combines multivariate pattern analysis (MVPA) with fMRI-based neuro-feedback to investigate complex brain activity patterns and their links to specific cognitive processes [1]. fMRI-based DecNef studies typically proceed in multiple stages. First, fMRI activity is recorded while participants perform a task. A machine learning classifier is trained on the distributed voxel activities to identify patterns associated with a specific cognitive state or behavior. These patterns become the target for neurofeedback training. During DecNef training, participants receive real-time visual feedback (i.e. of varying sizes of a disk) reflecting the likelihood that the current brain state matches the target, based on the trained classifier, and thereby increase in the monetary reward. A high proportion of participants fail at successfully self-regulating their brain activity in order to meet the target state during DecNef training and also in other neurofeedback training procedures, posing significant challenges in basic and clinical fields. The lack of intuitive access to one’s own brain activity can make DecNef training challenging for participants. Further, an important contributor to the difficulties encountered by participants may relate to differences in distributions between the training data used to construct the decoder and the data generated during neurofeedback test sessions (covariate shift—differences in feature space density between the perceptual training set and the DecNef testing set). When constructing decoders for neurofeedback, machine learning algorithms are typically trained on a dataset collected on a separate day employing different experimental conditions (i.e. task, stimuli) from those employed during neurofeedback training. Furthermore, fMRI activity can vary widely moment to moment due to several factors associated with neural and non-neural noise. Neural noise refers to variability in the fMRI signal that arises from spontaneous or task-unrelated fluctuations in neural activity. This includes trial-to-trial variability in neural activity, ongoing intrinsic brain dynamics, and state-dependent changes in attention, arousal, or cognitive strategy that are not directly driven by the experimental manipulations. Non-neural noise refers to variability in the fMRI signal that does not originate from neural activity. This includes physiological artifacts (e.g., cardiac and respiratory fluctuations), head motion, scanner-related instabilities (e.g., gradient drift or thermal noise), and other measurement-related factors that contaminate the BOLD signal. Due to these sources of noise, the decoder may not accurately capture the full range of variability in brain signals during DecNef training.

The aim of the present study is to develop a co-adaptation approach to improve the performance of brain decoders during neurofeedback training. The key idea behind co-adaptation is that not only the participants try to learn how to improve the induction of the target neural pattern, but critically that the decoder also learns from the participants’ attempts to induce the target state and adapt in a way that facilitates performance in subsequent attempts, while remaining more resistant to sources of neural and non-neural noise. The procedure thus involves an adaptive decoder algorithm that is updated in real time based on its predictions across neurofeedback trials. Co-adaptive machine learning algorithms were used previously in various studies in different fields, most importantly in the domain of brain computer interfaces (BCIs), robotics and autonomous systems. As detailed in the work of [2], most robotics and adaptive assistive technologies relied on the concept of reinforcement learning (RL) [3] to refine the system behaviors in response to changing user input or environments. Although a very important and effective technique, RL is impractical for the field of brain decoding especially with fMRI data, since the process of RL requires a large amount of data and iterations that does not conform with the experimental approach in fMRI DecNef studies. Thus, co-adapting machine learning models may offer a promising alternative. Previous studies in the field of BCIs using electroencephalogram (EEG) employed co-adaptation in different contexts. In the work of [4]–[6], co-adaptation was performed by using quadratic discriminant and linear discriminant classifiers that adapted to the user control of a computer interface through EEG. These approaches produced a significant improvement in the performance of the participants, especially since the EEG signal is known for random fluctuations and its non-stationary nature. The process of co-adaptation also eliminated the need for the calibration phase typically employed in the traditional BCI systems. The research of [7] demonstrated the use of Bayesian linear regression as a co-adaptation mechanism for a brain-machine interfaces through an invasive microwave implant. This method proved to improve the accuracy and robustness of the whole system through the continuous parameters fine-tuning which makes the framework able to adapt to both sudden and gradual changes through a periodical update to the decoder. [8], [9] used a smooth-batch linear discriminant classifier in an adaptive process over batches of samples for BCI training to minimize the effects of brain signal nonstationarities in EEG extractions of the power density spectrum over the alpha and beta bands. [10] proposed an adaptive system that calibrates and handles the nonstationarities of EEG data through an objective function that takes into account both performance and participant’s engagement. The framework tracks over time the user’s cognitive states, mood, and skill level to adjust the feedback for a more optimized learning process and sustainable engagement. [11] explored the role of co-adaptation in magnetoencephalography data. The study introduced a system with two convolutional neural network (CNN) architectures designed to improve classification performance and generalization over participants in BCI applications. The system is trained on pooled multi-subject data and includes an adaptive layer that fine-tunes for individual subjects in real-time at each prediction interval. This process allows the decoder to recalibrate based on the new data and mitigates the drawback of non-stationarity in MEG data. The role of co-adaptation has also been explored in functional MRI studies. [12] applied co-adaptive learning in BCI settings, such as fMRI and EEG, for real-time pain management. The protocol used an adaptive algorithm to reduce the intensity of pain stimulation delivered to participants based on decoding brain responses to previous pain stimuli. [13] implemented an unsupervised domain adaptation (UDA) on fMRI data to minimize the effect of cross-subject variability and data distribution shifts. The system is composed of a 3D CNN architecture to extract the spatiotemporal features and a UDA that calibrates the data distribution across subjects. This framework presented promising results that unfortunately was accompanied with some limitations due to the 3D CNN and domain adaptation resource intensive process that restrict potential real-time applications in fMRI contexts. Here, we present our novel co-adaptation approach for DecNef training in fMRI data, which manages to update in real time the decoder by retraining it based on the predictions produced during the neurofeedback trials. Initially, we performed simulations of the co-adaptation approach using an existing DecNef dataset [14]. Then, to validate the approach, we collected real time DecNef data from four participants and demonstrated that personalized decoders through co-adaptation can improve participants performance during DecNef training.

## II. SIMULATIONS OF THE ROLE OF COADAPTATION IN DECNEF TRAINING

### A. Methods

#### 1) Dataset and Experimental procedure

We modeled the role of co-adaptation during DecNef training by using the fMRI dataset from [14], which includes 18 participants. Please refer to the original publication [14] for additional details on the experimental procedures, scanner and imaging parameters. The dataset composed by three blocks of data:

- **The perceptual data** was used to train a decoder to classify the category of the visual images presented to the participants during fMRI scanning, namely, living vs non-living (i.e. dogs vs scissors). The data is divided into 4 different runs with each run containing approximately 209 or 210 volumes. After the preprocessing (detrending, z-scoring and volume realignment), applying a mask of the fusiform cortex over the volumes and the extraction of fMRI volumes of interest for decoding, the number volumes ranged between 360 and 386 per participant.
- **The neurofeedback data** was used for testing the decoder during DecNef training. It was also partially used for the additional step training phase of the co-adaptation and the testing phase of the co-adaptation evaluation as well. The data was gathered in 4 sessions performed on different days, containing 8 runs for initial 3 sessions and 2 runs for the last session. FMRI data were collected while participants attempted to induce the target brain pattern by changing their mental strategies, which unbeknownst to them, was associated with the living class (dog). Participants received feedback in the form of a feedback disk, and monetary reward contingent on the size of the feedback disk which was based on the probability of the decoder regarding the living class.
- **The imagery data** was collected while the participants attempted to simulate, via mental imagery, living and non-living stimuli presented previously. These data were used to test the generalization of the co-adaptation process. The imagery data were collected in 2 runs of 206 volumes each.

As summarized in Figure 1. First, a decoder is trained on the perceptual data to classify the living and non-living nature of the perceptual stimulus (see [14] for more description on the data collection).

**Fig. 1.**
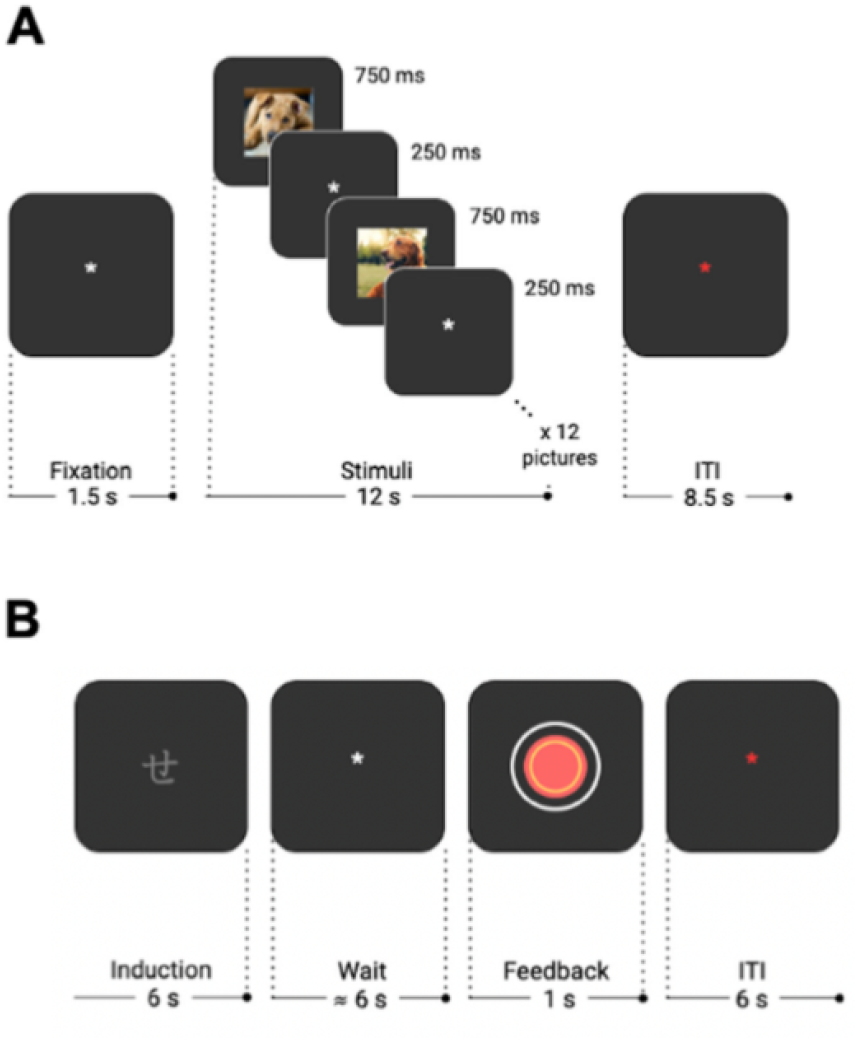
(A) Illustration of the experimental procedure used during the perception phase for subsequent decoder construction and (B) example of a DecNef training trial in which participants were required to induce a brain pattern that maximizes the size of the feedback disk, which unbeknownst to the participants, corresponded to the probability of the decoder regarding the living class.

Imagery data on the dogs and scissors were also collected. Then, the DecNef training procedure was conducted. The purpose of this experiment was to train participants to induce training and also for the neurofeedback training sessions.brain activity patterns similar to those of the perceptual living class, without awareness of the contents that those patterns represent. In the present study, we developed a reliable decoder through a co-adaptation procedure, a decoder that can generalize between the different experimental contexts and data distributions. We also aimed to develop a decoder that can be trained as fast as possible using the minimum amount of resources, since the collection of fMRI data typically produces relatively small amounts of data for machine learning models in a single session.

#### 2) Decoder construction based on the perceptual data

Previous DecNef studies typically used the logistic regression classifier as the main decoder for the experiment. In our case, we decided to test various machine learning models from diverse algorithmic families before conducting our analysis. These algorithms included tree based models, ensemble based models, instance based models, bayesian models and linear models. This inventory of models that we selected consists of: ExtraTrees(ET), the k-nearest neighbors (KNN), RandomForest (RF), Gradient Boosting (GB), Support Vector Machine (SVM), Logistic Regression (LR), DecisionTree (DT), Multilayer Perceptron (MLP) and Naive Bayes (NB). Since the neurofeedback process depends on the prediction of the model to provide the feedback signal to the participant, we decided to rely on the F1 score metric, also known as balanced accuracy (see equation (1)) to evaluate our models both for the decoder training and also for the neurofeedback training sessions. The F1 score metric represents a reliable tool to evaluate the performance of the coadaptation process as explained in the work of [15], since the neurofeedback data can be considered data from a single class given the participant’s requirement to induce a single target representation (i.e. dogs) and is therefore a highly imbalanced dataset.

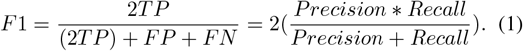

Where TP is True Positive (the model correctly identifies the outcomes as positive), FN is False Negative (the model incorrectly identifies the outcomes as negative) and FP is False Positive (the model incorrectly identifies the outcomes as positive) with the latter non-existent in the case of testing on the neurofeedback data including only one class. The F1 score metric during decoder construction was validated using the leave-one-run-out cross-validation procedure with the grouping depending on the runs available for the training data in the perceptual sessions. More than half of the participants (10 participants) had a better F1-score in the cross validation test with the use of the Logistic Regression classifier (the overall mean across the 18 participants was 86.5%). Additionally, 3 participants had better results with the RandomForest classifier (the overall mean across the 18 participants was 83.7%). Based on these results we decided to select the logistic regression classifier as our default decoder in the rest of our experimentation with the co-adaptation, which is also convenient for comparing our proposed methodology to previous studies.

#### 3) Co-adaptation protocol

Decoded neurofeedback training was performed using our Python-based pipeline [16]. The co-adaptation protocol used in the initial simulations that were performed on our prior data [14] involved two phases. First, we trained a decoder on perceptual stimuli data to classify living and non-living categories. Then, the decoder was applied to the neurofeedback data collected while participants attempted to induce the target pattern (“living class”). Based on the classifier’s predictions, the model was retrained on the fly using neurofeedback examples that passed a confidence threshold. Specifically, we selected the neurofeedback training examples based on the probability of decoding the target class. We applied two confidence thresholds—65% and 80%—in separate simulations. The rationale for testing these two thresholds was to examine the trade-off between data quantity and quality in the co-adaptation process. A lower threshold (65%) provided the decoder with more training samples but at the cost of increased noise, whereas a higher threshold (80%) likely ensured better representations of the target class but fewer examples for the decoder to adapt to.

As described above, fMRI volumes for which the model’s predictions surpass the confidence threshold are sequentially and cumulatively used to retrain the decoder. Specifically, these selected examples are concatenated, and their labels are reassigned to match the target class before being incorporated into the training set. Each time an fMRI volume exceeds the confidence threshold, the volume is selected for co-adaptation. The retraining of the decoder always takes place after the relevant fMRI volumes corresponding to the induction period (accounting for the hemodynamic delay) have been evaluated. The co-adapted decoder is then tested on the examples that do not pass the confidence threshold and the decoder predictions are then used to evaluate the decoder’s performance at each iteration. This retraining process remains computationally efficient due to the use of machine learning algorithms that require minimal parameter updates per iteration. In contrast, deep neural networks, which involve extensive weight adjustments, demand significantly more time and resources, making them less practical in this context. Additionally, neurofeedback experiments inherently produce limited fMRI data, as participant scanning time is constrained. Given these factors, the co-adaptation protocol employed here offers an optimal balance between adaptability and efficiency, outperforming reinforcement learning and deep neural networks for real-time neurofeedback applications.

### B. Results

Below, we present the results from multiple simulations. First, we examine the effect of the co-adaptation protocol on the neurofeedback data using two different confidence thresholds (see Methods). Next, we replicate this analysis while balancing the amount of training and test data to ensure that the observed effects of co-adaptation are not merely due to an increased number of training samples or decoder biases associated with this factor. We also validated the stability of the co-adapted decoder over the course of the neurofeedback sessions by means of a drift analysis. We assessed the drift by testing the model at each step of the co-adaptation on a left-out run of the perceptual data. In other words, the perceptual decoder was trained using three of the four perceptual runs and then, at every step of the coadaptation with the neurofeedback data, the coadapted decoder was tested to discriminate the perceptual classes of the left out run not used for decoder training. This procedure was repeated in a cross-validation manner using the leave-one-run out method, and the F1-scores were averaged across runs to ensure more reliable estimates.

#### 1) Co-adaptation effects on neurofeedback training

The volumes that passed the decoder’s confidence threshold of 80% for the living class ranged between 411 and 892 samples depending on the participants with a mean of 636.35 number of volumes and standard deviation of 122.06 per participant. We note that these samples are excluded from the testing dataset before we start the co-adaptation procedure, and also in the baseline assessment in which the decoder was not co-adapted. When the decoder was trained on the perceptual data then tested on the whole neurofeedback data (i.e. including all the examples regardless of whether or not they surpass the confidence threshold), we observed an average F1-score of 73.96%. In the co-adaptation test, we initially trained a decoder on the perception data and then we iteratively re-trained the model during co-adaptation using the neurofeedback examples obtained in sequential order that passed the confidence threshold (see II-A). This co-adapted decoder was compared to another decoder trained only on the perception data. Notably, both decoders were tested on the same neurofeedback volumes that did not pass the confidence threshold in order to prevent any bias or advantage in the co-adapted decoder resulting from being trained and tested on the same neurofeedback examples. Figure 2 (A) shows the evolution of the average F1-score for all the participants using the co-adapted decoder, demonstrating how it significantly surpasses the average F1-score of the non-coadapted decoder. Additionally, a drift test was conducted to see if the decoder performance on the training data degraded during the co-adaptation procedure. This drift test witnessed a negligible decay from 86.63% at the beginning to 85.36% average F1-score at the end of the coadaptation process for a cross-validation test that left one of the training runs out each time (see Figure 2 (A), red dotted lines). The outputs probability distribution of the co-adapted decoder also confirms the improvement in the performance compared to the original decoder as shown in Figure 6 (In the appendix). Figure 2 (B) illustrates the average F1-score across all participants for the co-adapted and non-coadapted decoder. The boost in the performance of the co-adapted relative to the non-coadapted decoder is significant (t(17) = 24.11, p < 0.0001).

**Fig. 2.**
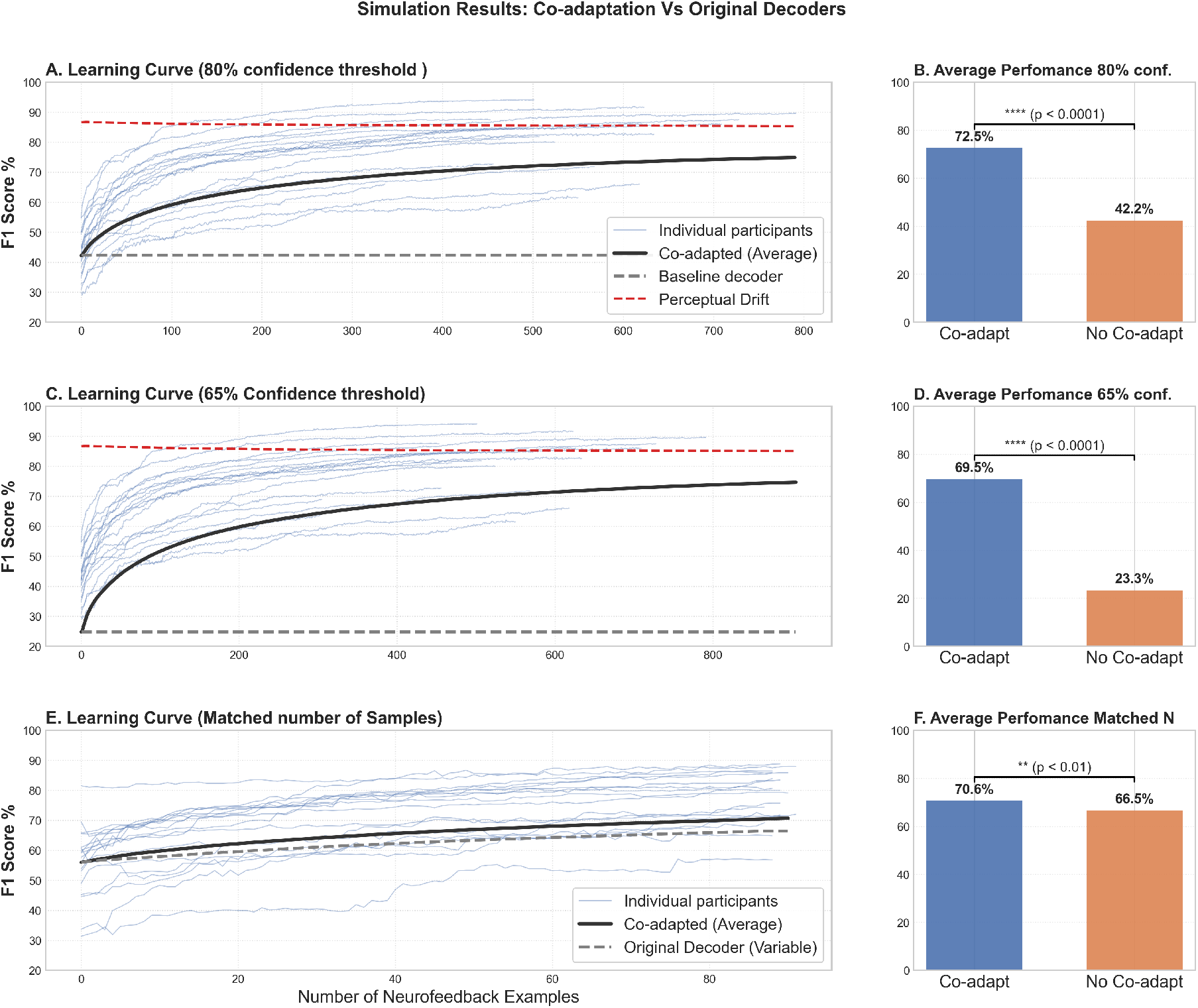
The average F1-score of all participants for the co-adapted decoder with 80% confidence threshold compared to the non-coadapted decoder tested on the same number of neurofeedback samples. The coloured lines illustrate the performance of individual participants.

We repeated the simulations by re-training the co-adapted decoder using neurofeedback examples that passed the 65% confidence threshold. Here, the impact of co-adaptation may be higher due to the use of more neurofeedback examples. Figure 2 (C) illustrates the results. Likewise, the drift test showed that the co-adapted decoder performed well in the left-out set of the perceptual data at every step of the coadaptation, with little variation from 86.63% to 85.09% average F1-score in cross-validation (see Figure 2 (C), red dotted lines). Performance of the co-adapted decoder was significantly higher (t(17) = 26.34, p < 0.0001), as shown in Figure 2 (D).

It might be argued that the performance gains observed with decoder co-adaptation could arise from a trivial linear bias, whereby all decoder outputs are uniformly shifted upward rather than reflecting a meaningful refinement of the decision boundary. To address this possibility, we compared the distributions of decoder probability outputs produced by the original and co-adapted decoders across the full dataset (see Fig. 6 in the Appendix). If co-adaptation simply introduced a global bias, one would expect a uniform increase across all probability values. Instead, the co-adapted decoder exhibited a non-linear redistribution of output probabilities, characterized by selective increases for trials exceeding the 50% decision threshold and relatively minimal changes for sub-threshold trials. This pattern indicates that co-adaptation does not act as a global gain term, but rather sharpens the decoder’s decision boundary to better distinguish target from non-target states.

#### 2) Co-adaptation using the same quantity of data within both training and testing

It might be argued that the increased performance of the co-adapted decoder is due to the higher amount of training data relative to the non co-adapted decoder. Even though this argument is somehow irrelevant to the goal of improving the DecNef training performance, we tested the procedure by matching the number of samples used for training and testing both the co-adapted and non-coadapted decoder. We trained both decoders partially on the perceptual data. Then, we continued the training sequentially during co-adaptation on the neurofeedback data. Likewise, in the case of the non-coadapted decoder, we continued the training but here we used the remaining perceptual data. The initial decoder trained on the perceptual data was initialised using all the examples from the non-living class (scissors) and half the examples of the target class (dog). Then, on each step of the retraining process, an example from the neurofeedback data was added to the training data of the co-adapted decoder, while in the case of the non-coadapted decoder an example from the living class of perceptual data was added on each step of the retraining process, thereby keeping an equal number of volumes in training and test sets of both co-adapted and non-coadapted decoders. The replacement process involved only examples of the target class, since this is the only relevant class for the neurofeedback training. We conducted this test using the 80% confidence threshold since the number of samples that passed this threshold are even more than the number of samples from the living class in the perceptual data, which ensures a better quality data with no compromise on the data quantity. It is worth noting that in this test, the amount of data for decoder construction and for testing was much reduced compared to the previous tests, which can explain why the benefit of co-adaptive training illustrated in Figure 2 (E) and Figure 2 (F) was relatively lower by comparison, but still significantly higher compared to the non-coadapted decoder (t(17) = 3.33, p < 0.01).

#### 3) Cross-domain co-adaptation from DecNef to mental imagery

In order to further test the generalization ability of the co-adaptation procedure, we validate using the imagery stimulus data in the testing phase. The imagery data contained fMRI examples when participants were simulating via mental imagery the properties of the living and non-living stimulus class. Likewise, during DecNef participants must internally generate a representation that resembles the target state, even if they are unaware of the specific content being represented (i.e. dogs). Therefore, we reasoned that co-adapting the perceptual decoder with the neurofeedback examples that meet the confidence threshold for the target class, ought to improve decoding performance of the contents represented in mental imagery. It is also important to note that in this cross-domain test, the co-adapted decoder will not have the advantage of being re-trained and tested on (neurofeedback) data with a similar distribution, thereby, preventing any biases in the computation of decoder performance. Furthermore, here the co-adapted decoder is tested on examples belonging to two classes (living vs non-living), thereby, preventing any biases in the computation of decoder performance. First, the decoder was trained on the perceptual data. Then, the decoder was co-adapted using the neurofeedback examples according to the chosen confidence threshfolds (65% or 80%). Finally, we test both the original decoder and the co-adapted decoder at each step of co-adaptation on the imagery stimulus data. Figure 3 shows the average F1-score for the different decoders with and without co-adaptation. Co-adaptation led to improved decoding performance relative to the non co-adapted decoder with both the with 80% confidence threshold (t(17) = 4.52, p < 0.001) and the 65% confidence threshold (t(17) = 4.57, p < 0.001). The improvement was higher than 2 - 3 points, which is remarkable given the cross-domain nature of the problem and considering also the small size of the test data. The slight reduction in the size of the improvement for the co-adapted decoder with the 80% confidence threshold compared to the 65% confidence threshold may be due to the lower size of the data used for co-adaptation.

**Fig. 3.**
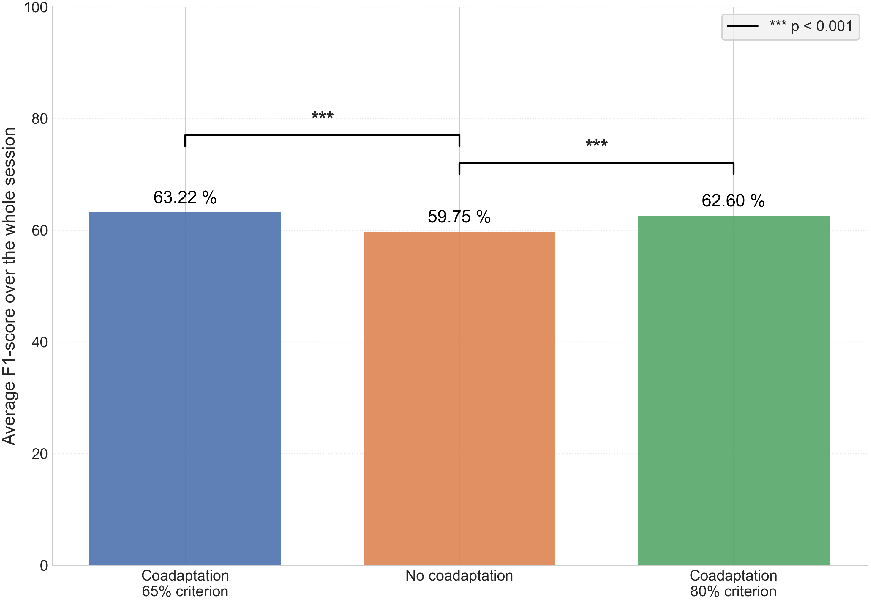
Average F1-score for the co-adapted decoder with both 65% and 80% confidence threshold compared to the original perceptual decoder tested on imagery stimulus data.

Following the simulations of the co-adaptation procedure using our existing dataset, we then performed a closed-loop DecNef training experiment with four participants to provide proof-or-principle evidence of the effect of co-adaptation on a real-time setting. This test will further allow us to demonstrate whether co-adaptive training can have consistent performance improvements that can be visualized within single participants.

## III. Testing the role of co-adaptation in real-time closed-loop neurofeedback training (Proof of principle)

### A. Methods

#### 1) Participants

We recruited four participants (3 females and 1 male, aged between 22 and 25 years). The experiment was approved by the BCBL Ethics Board and conducted in accordance with the Declaration of Helsinki. All participants provided written informed consent. At the end of the experiment, participants received monetary compensation at a rate of 15 C per hour.

#### 2) MRI settings

Whole-brain data were acquired using a 3T Siemens Magnetom Prisma-fit whole-body MRI scanner with a 64-channel head coil at the BCBL facilities. A gradient-echo echo-planar imaging sequence was employed with the following parameters: repetition time (TR) = 2000 ms, echo time (TE) = 28 ms, flip angle (FA) = 74°, field of view (FoV) = 192 mm, and 34 axial slices with no inter-slice gap, with a voxel resolution of 3 × 3 × 3.5 mm^3^. At the start of each fMRI run, five volumes were acquired to allow for MRI heat-up and ensure steady-state magnetization; these volumes were excluded from the analysis. Visual stimuli were presented on an out-of-bore screen, viewed by participants through a mirror attached to the head coil. To minimize head movements, participants were instructed to keep as still as possible, and foam padding was used to cushion the head coil. They were also instructed to avoid respiration changes. Participants wore padded headphones to reduce scanner noise, facilitate auditory stimulus presentation, and enable communication with experimenters via a compatible microphone. At the end of each run, participants were offered a short break of 1-2 minutes. A high-resolution T1-weighted structural MPRAGE scan was acquired: TR = 2530 ms, TE = 2.36 ms, FA = 7°, FoV = 256 mm, 1 mm isotropic voxel resolution, 176 slices.

#### 3) fMRI Scans for Decoder construction

The model construction session involved four perceptual runs. During these runs, participants were presented with pictures of ‘Dogs’ and ‘Scissors’ from the THINGS database [17]). Each perceptual run consisted of seven ‘Dogs’ and seven ‘Scissors’ trials, lasting approximately 8 minutes. A trial began with a 1500 ms white fixation on a black background, followed by the sequential presentation of 12 example pictures from a specific visual category (‘Dogs’ or ‘Scissors’) for 750 ms each. The picture presentations were interleaved with 250 ms visual fixation periods. This block design was used to improve the signal-to-noise ratio for specific category representations (see Figure 1). The same images were used for each category across trials but were randomly presented in each mini-block of 12 images. To model the brain activity patterns associated with each category and enhance the separation of these patterns, an 8500 ms inter-trial interval (ITI) with a red fixation point at the center of the screen was included. Throughout the perceptual runs, participants were instructed to focus on the pictures and actively engage with their meaning.

#### 4) FMRI volumes preprocessing

The raw fMRI images were processed through a customized pipeline using AFNI v21.0.05 [18], which was implemented via the Nipype v.1.4.1 [19] interface for Python 3.6. A similar preprocessing was done for the decoder construction dataset and real-time processing during DecNef. The first experimental volume acquired after the heat-up volumes in the model construction session was used as the functional reference for each participant. This reference DICOM file was converted to NIfTI format using Dcm2Niix [20] and de-obliqued to align with cardinal coordinates using the AFNI 3dWarp command. Subsequent volumes were co-registered to the reference volume using the AFNI 3dvolreg command with heptic (7th order) Lagrange polynomial interpolation [21]. Task logs from OpenSesame were used to label all fMRI volumes, identifying stimulus categories, trial onset times, and corresponding runs. To prepare the data for decoder training, we linearly detrended the fMRI time series to remove drift in each run. Here we elected to use scans between 4 and 6 seconds after the onset of the first image presentation —comprising approximately 1 or 2 volumes per trial— were selected. This means that the amount of training data in this co-adaptation test was reduced to 35 examples per class, in order to avoid imbalance in the number of examples used in the decoder training session and the subsequently neurofeedback testing session with co-adaptation. This consideration is also relevant for devising experimental procedures that include both decoder construction and DecNef-based co-adaptation within the same experimental session, since participants’ time spent on the MRI scanner within a single session is limited. Importantly, we ensured that the performance of the decoder was high enough to proceed with the decoded neurofeedback training. These selected perceptual samples were then stacked and z-scored by subtracting the mean value at each voxel and dividing by the standard deviation. he data for training the decoder (and also for DecNef) was based on a region of interest in the visual cortex. We create a mask in standard MNI space comprising the following areas: lingual gyrus, occipital fusiform and temporo-occipital fusiform. Then, this mask was registered into the native functional space on each participant using a linear transformation using FSL FLIRT.

#### 5) Algorithm for decoding and co-adaptation

In order to select the decoder for neurofeedback training, we tested several decoders using a leave-one-run out cross-validation procedure on the perceptual data. Similar to the case of the simulated data, the logisitic regression decoder scored the best results with an average F1-score of 82.02% across the 4 participants. Consequently, a logistic regression decoder was trained on the perceptual data to predict the living and non-living classes. Then, we use the pre-trained model to make predictions on the DecNef session. Here, for each example in which the decoder predictions pass the 65% confidence threshold, the corresponding volumes are added in real time to a newly created training dataset that includes the perceptual data. These volumes are labeled with the target class. A new training process of an uninitialized decoder is performed in this dataset. Co-adaptation was performed once the relevant volumes associated with induction related activity on a given trial were evaluated. This co-adapted decoder is then saved and used to make predictions in the next trial. The same process is iteratively performed for new volumes that pass the confidence threshold. The sequence of events on an example trial were similar to those depicted in Figure 1b, except that the feedback period was increased from 1 to 1.5 seconds and the ITI was also increased from 6 to 10 seconds, in order to further mitigate any carryover effects from previous trials.

#### 6) DecNef training

Following the decoder construction session, participants came back for another session of DecNef on a different day. Similarly to our prior study, participants were asked to induce a brain pattern that maximizes the size of the feedback disk bigger, but they were kept unaware of the specific content associated with the feedback (Figure 1b). Participants were also instructed to change their brain activity through mental strategies and strictly avoid movement or respiration changes, in order to maximize the size of the feedback disk. In order to test the effect of the co-adaptation, we alternated between fMRI runs in which no co-adaptation was used and runs in which co-adaptation was performed. Specifically, DecNef training began with two runs in which no-coadaptation was used. This was done to use the examples that surpassed the confidence threshold in these two initial runs for re-training the classifier in order to maximise the effect of co-adaptation in the third run. Subsequently, the fourth run of neurofeedback training switched back to the original decoder which had not been co-adapted. Finally, the last fifth run of training again employed co-adaptation, and the decoder was further re-trained using the examples that passed the confidence threshold regarding the living class on the fourth run. Also, the decoder continued to be co-adapted on a trial by trial basis using the examples that passed the confidence threshold on a given co-adaptation run. We also note that due to an experimenter error, the first run of participant 2 included co-adaptation but this was again switched to no co-adaptation in the second run.

### B. Results

Figure 4 illustrate the performance of the decoder during neurofeedback training for each of our 4 participants throughout the different runs, which alternated between non-coadapted and co-adapted decoders (see III-A). The results shown in this Figure illustrate that the likelihood of inducing the target neural pattern during DecNef was higher for runs in which co-adaptation was used relative to a non-coadapted decoder. This performance enhancement is seen on every single participant tested. Table I reveals the F1-score of the 4 participants showing that the average F1-score for the participants was higher during co-adaptation compared to no co-adaptation runs (t(3)=3.43, p < 0.05).

**TABLE I.**
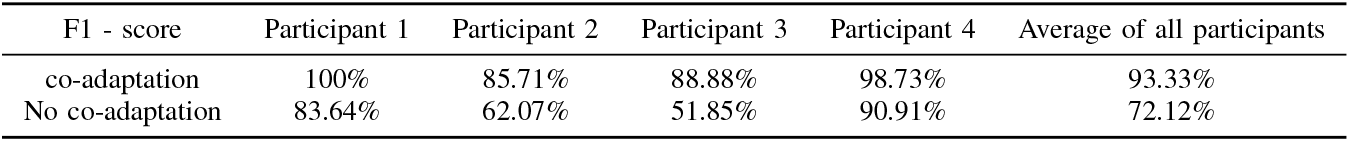
F1 score results of the 4 participants with co-adapatation compared to the control decoder (original decoder Non co-adaptated)

| F1 - score | Participant 1 | Participant 2 | Participant 3 | Participant 4 | Average of all participants |
| --- | --- | --- | --- | --- | --- |
| co-adaptation | 100% | 85.71% | 88.88% | 98.73% | 93.33% |
| No co-adaptation | 83.64% | 62.07% | 51.85% | 90.91% | 72.12% |

**Fig. 4.**
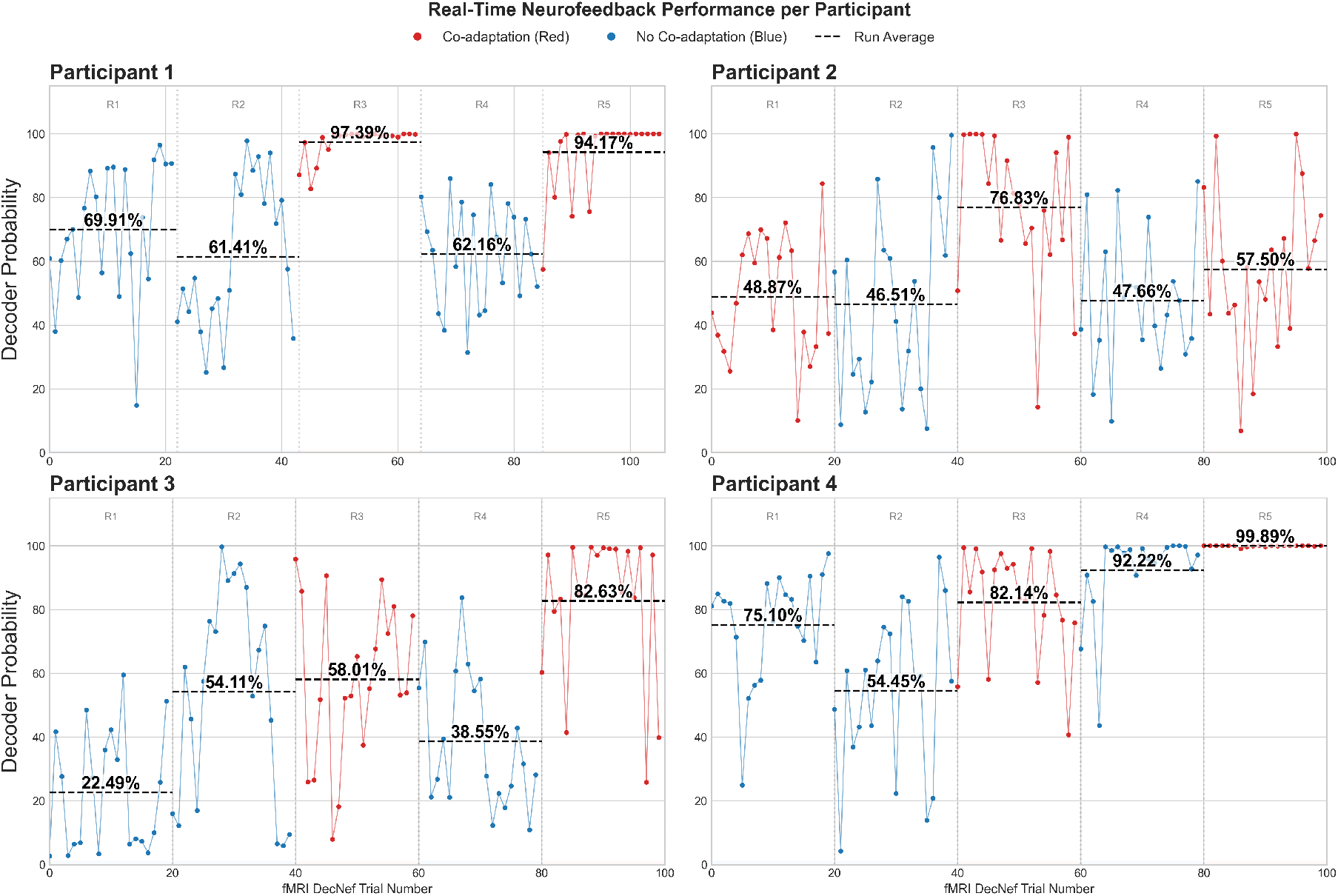
Illustration of decoder predictions for the participants regarding the living class with co-adaptation (red) and without it (blue). Note that here each of the fMRI trials in DecNef may have 2-3 examples. (A) participant 1, (B) participant 2, (C) participant 3 and (D) participant 4.

Lastly, we also carried out the same experiment but using the non-living (scissor) as the target class to be induced during DecNef. Thereby, two of the four participants came back for a final session after they had performed the experiment with the living class. This allowed us to further test the generalization ability of the co-adaptation process. As can be seen in Figure 5, the same improvement in decoding performance during neurofeedback training was observed in each of the two participants tested.

**Fig. 5.**
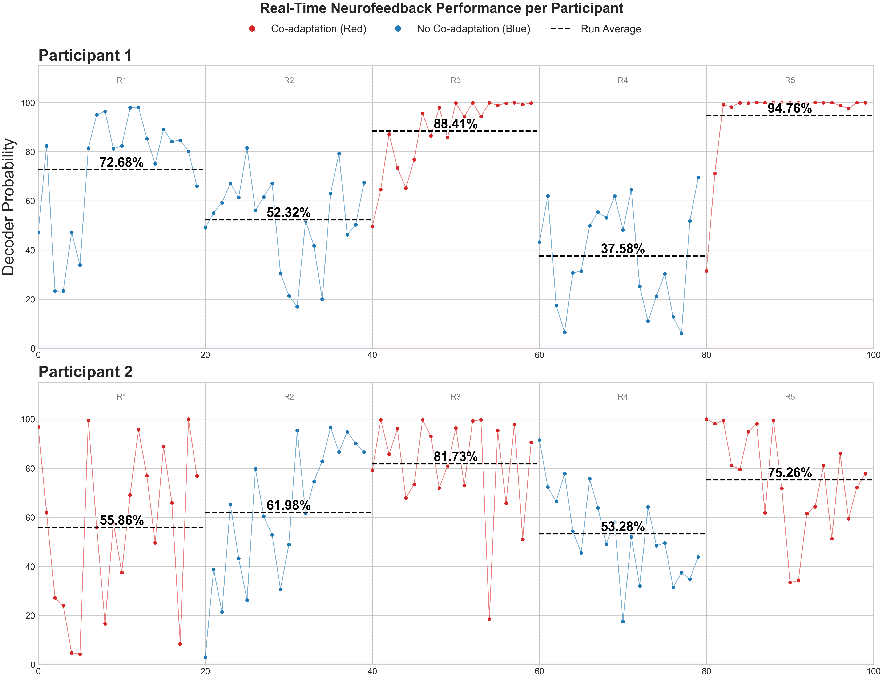
Illustration of decoder predictions for the participants regarding the non-living class with co-adaptation (red) and without it (blue). Note that here each of the fMRI trials in DecNef may have 2-3 examples. (A) participant 1 and (B) participant2.

**Fig. 6.**
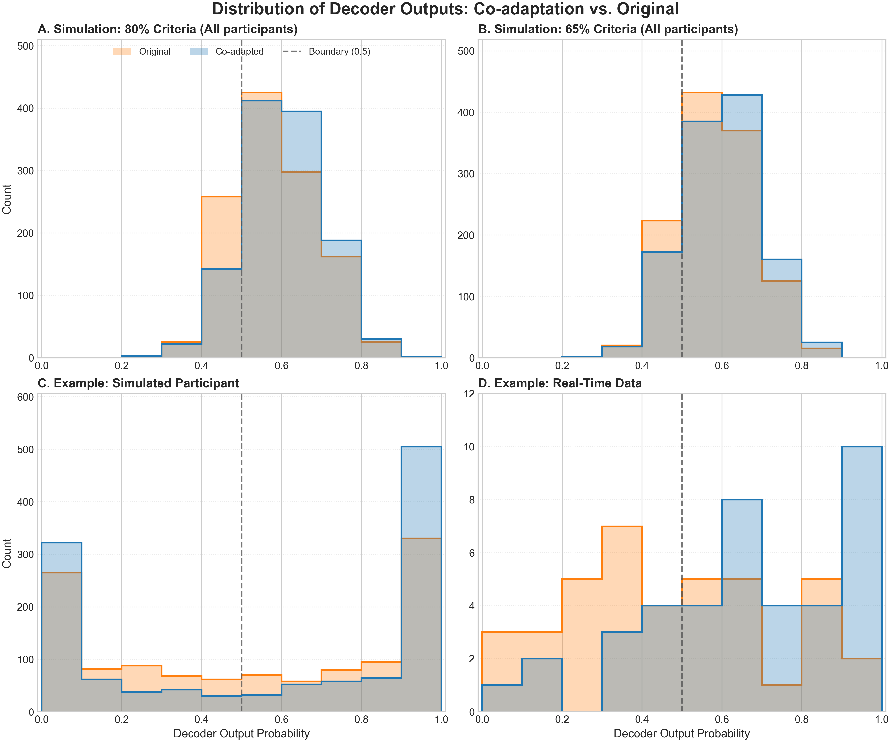
Distribution of decoder output probabilities for the original (orange) and co-adapted (blue) decoders. (A) Histogram of output counts per average probability bin for all participants in the simulated data with 80% confidence threshold. (B) Histogram of output counts per average probability bin for all participants in the simulated data with 65% confidence threshold. (C) Histogram of output counts per probability bin for an individual participant of the simulated data. (D) Histogram of output counts per probability bin for an individual participant of the real-time data.

## IV. Discussion

The aim of this study was to develop a co-adaptation approach to enhance brain decoder performance during neuro-feedback training using fMRI. Unlike conventional neurofeedback, where participants alone try to induce the target brain state, co-adaptation allows the decoder to dynamically learn from participants’ attempts and update its predictions in real time. To test this approach, we first conducted simulations using an existing DecNef dataset [14] and then performed a proof of principle real-time co-adaptive neurofeedback training with four participants. This co-adaptive approach is consistent with previous work in EEG-based BCIs, such as that of [4], which demonstrated significant performance improvement in sensory-motor neurofeedback tasks using motor imagery, through an adaptive machine learning procedure based on linear discriminant analysis. However, in prior work using adaptive neurofeedback protocols [8], [9] participants were consciously aware of the relevant content associated with the target state of neurofeedback training (e.g. type of motor imagery task), while in fMRI DecNef studies participants are typically unaware of the target of neurofeedback training.

Our simulation results demonstrated a significant improvement in decoder performance due to co-adaptation, both with relaxed and stringent confidence thresholds for selecting training examples for co-adaptation. Compared to a static decoder trained solely on perceptual data, a decoder co-adapted with neurofeedback data achieved higher F1-scores, confirming its ability to refine classification accuracy dynamically. This effect persisted even when reducing the size of the training set and, importantly, the benefits of co-adaptation cannot be attributed to extraneous factors such as classifier bias during retraining, given that the use of balanced datasets consistently led to improvements compared to non-co-adapted decoders. To assess the generalizability of co-adaptation, we cross-validated the perceptual decoder co-adapted with neurofeedback data by testing it on mental imagery data. We observed reliable improvements of the co-adapted decoder when tested on the imagery data, despite the absence of retraining on imagery data. This cross-domain generalization test highlights the robustness of co-adaptation in our simulations. Moreover, the drift analysis highlights the stability of the co-adapted decoder throughout the neurofeedback training process. Performance remained consistent across co-adaptation steps, indicating that the decoder preserved its ability to generalize to unseen perceptual data despite ongoing updates. Importantly, there was no evidence of performance degradation on the training data, suggesting that the co-adaptation process did not lead to overfitting or biases. These findings support the robustness of our approach and reinforce its suitability for sustained use in neurofeedback applications.

As a proof-of-concept demonstration, we examined the effect of co-adaptation in a real-time DecNef training experiment. Alternating between co-adapted and non-co-adapted fMRI runs, we found consistent improvements in participants’ ability to induce the target state. Additionally, two participants repeated the experiment with a different target class (non-living objects), and the same performance enhancement was observed, indicating that the co-adaptation effect can generalize when a switch in the target representation is introduced. Nevertheless, these results should be interpreted with caution given the small sample size (n = 4). The real-time experiment serves as a proof-of-concept. While the effect of co-adaptive training was evident within each participant, the low sample size limits the statistical power and generalizability of the findings in the general population. These results should be interpreted as preliminary evidence warranting larger-cohort validation.

It might be argued that the signal driving decoder co-adaptation during decoded neurofeedback does not reflect selective reinforcement of the target representation (dogs), but instead arises from stimulus-driven responses to the hiragana character repeatedly presented during induction. Given that the fusiform gyrus is responsive to letters and symbols, it is conceivable that the decoder could gradually adapt to features associated with hiragana viewing, potentially transforming into a classifier that distinguishes dogs + hiragana from scissors rather than dogs per se. While this is an important consideration, several lines of evidence argue against this interpretation. First, prior work using the same DecNef framework demonstrated selective modulation of the neural representation of dogs accompanied by corresponding behavioral effects on dog search performance [14], indicating that training reinforces category-specific representations rather than stimulus-generic visual responses. Second, the decoder used here was trained on dog and scissors images and achieved high classification accuracy, making it unlikely that univariate fusiform responses to the hiragana symbol would dominate the feedback signal. Third, if co-adaptation primarily reflected a shift toward a dogs + hiragana versus scissors decision rule, one would expect a substantial degradation in decoder performance on the original perceptual localizer data. Instead, we observed only a small (1–2%) reduction in accuracy, consistent with fine-scale adjustments around an already well-established dog–scissors boundary rather than a qualitative re-encoding driven by hiragana responses. Together, these considerations support the interpretation that co-adaptation refines task-relevant category representations, rather than exploiting incidental stimulus-evoked activity.

A potential concern regarding the real-time DecNef experiment is the interleaved run structure, particularly in light of an experimenter error affecting Participant 2, for whom the first run involved co-adaptation rather than the intended no–co-adaptation condition. However, any learning acquired during this initial co-adaptation run would be expected to carry over and elevate performance in the subsequent, second no–co-adaptation run. Importantly, such carryover would reduce the contrast between conditions and therefore work against, rather than in favor of, our hypothesis. Despite this conservative bias, a reliable advantage for the co-adaptation condition was still observed. More generally, the interleaved within-subject design was deliberately chosen to provide a more stringent test of co-adaptation effects. Separate-groups or session-wise baseline comparisons across different days are more vulnerable to non-stationarities that evolve over the course of 1 h fMRI sessions, including fatigue, learning, attentional fluctuations, and scanner drift. Interleaving co-adaptation and no-coadaptation conditions within a session minimizes these potential confounds and provides a more stringent test of co-adaptation effects. Finally, we note that an external baseline is provided by our previous study [14], in which participants performed the same DecNef induction task without decoder co-adaptation and showed lower overall performance. Taken together, these considerations support the robustness of the observed co-adaptation effects.

Standard DecNef training studies typically employ three sessions across different days. Prior fMRI DecNef studies do not often report significant learning curves during training, with some exceptions [14] [22]. Our findings suggest that co-adaptation could enhance the magnitude of DecNef effects on both neural and behavioral outcomes, potentially reducing the number of sessions required for effective modulation. Even in cases where participants fail to learn how to reliably induce the target brain pattern, DecNef training may still be effective through a reinforcement learning mechanism, since the mere induction of the target state—even if occurring in only half of the trials—could potentially lead to lasting neural and behavioral changes given that the brain state is paired with a reinforcer [23] [24]. However, it is clear that developing new DecNef procedures that enhance participants’ ability to induce the target state with greater precision and consistency could be transformative. By improving participants’ control over neural modulation, we can better evaluate the impact of DecNef on brain function, cognition, and behavior.

These results highlight the potential of co-adaptation to address key challenges in brain-computer interface (BCI) research and fMRI-based DecNef studies, particularly in mitigating individual variability and enhancing system adaptability. The iterative nature of the protocol allows for continuous refinement of the decoder, making it increasingly responsive to each user’s unique neural patterns over time. This adaptability is especially valuable for clinical populations, such as individuals with neurodevelopmental conditions, where variations in brain activity—due to neural and non-neural noise, motivation, or fatigue—often hinder performance. Co-adaptation may also enhance participant engagement and motivation during neuro-feedback training. By dynamically adjusting to the user’s brain activity, the system provides a sense of self-control, making the process more rewarding and intuitive. This could be particularly important in training protocols where inducing the target state is challenging. In our experience with DecNef studies, participants often reported that the strategy they had been using to maximize the feedback disk’s size suddenly stopped working, forcing them to explore new approaches. These disruptions could stem from non-neural noise, fluctuations in attention or motivation, as well as non-stationary effects caused by physiological noise or drifts/instability in the MRI scanner. The co-adaptation procedure may help mitigate these issues by enhancing the decoder’s robustness to non-stationary signals [8]. This not only allows for more accurate predictions of the intended neural patterns induced by participants but also ensures a more stable and personalized neurofeedback training experience.

A possible mechanism underlying these effects is that co-adaptive training facilitates learning by effectively reducing the search space faced by participants during induction. In conventional DecNef, participants must explore a high-dimensional and largely unconstrained neural space to discover strategies that elicit the target pattern, often without explicit awareness of the target representation. Co-adaptation may alleviate this challenge by progressively aligning the perceptual decoder with each participant’s idiosyncratic neural patterns expressed during successful induction attempts. In doing so, the decoder becomes increasingly sensitive to neural activity patterns that the participant can reliably produce, thereby constraining the feedback landscape and making effective strategies more discoverable. From this perspective, co-adaptation does not merely enhance classification accuracy, but also reshapes the mapping between neural activity and feedback in a way that supports participant learning. This interaction between decoder adaptation and participant exploration may help explain why co-adaptive DecNef leads to more consistent induction of the target state, even when explicit learning effects are typically weak or absent in standard DecNef paradigms.

While our findings illustrate the benefits of co-adaptation, some limitations remain. Performance gains were less pronounced in cross-context scenarios, such as when co-adaptation was applied to neurofeedback training data but tested on imagery data. This suggests that co-adaptation may require additional strategies—such as domain adaptation techniques [25]—to improve generalization across different experimental conditions. Additionally, we consider the issue of the stopping criteria during co-adaptive training. Co-adapting the classifier indefinitely might lead to biases in the classification predictions. However, our drift analyses showed that this was not the case. This drift analysis can be easily incorporated in the real time decoding pipeline to monitor potential drifts from the target representation.

To further advance co-adaptive neurofeedback protocols for personalized brain training, future research may explore unsupervised domain adaptation techniques, which could recalibrate the system based on similarities between target representations and neural patterns induced during neurofeedback [26]. Additionally, representational learning algorithms, such as autoencoders and contrastive learning, could be employed to identify robust neural representations for decoded neurofeedback training. These advancements could be facilitated by the use of more accessible and portable systems, such as fNIRS, which would allow for the collection of a larger number of training and testing examples and expand the practical applications. Co-adaptation protocols represent a promising advancement in DecNef training, offering a practical approach to enhancing decoder performance and participant engagement in both clinical and non-clinical settings. By making neurofeedback training more adaptive, personalized, and effective, this approach has the potential to accelerate neural modulation, improve learning outcomes, and expand the applicability of DecNef techniques in cognitive neuroscience and clinical interventions. The tools are made openly available to the scientific community to foster collaboration and advance decoded neurofeedback research.

## Open Science Statement

The Python library for coadaptive DecNef and the data can be found in the following github repository: https://github.com/pydecnef/PydecNef

## Appendix

